# BERTE: High-precision hierarchical classification of transposable elements by a transfer learning method with BERT pre-trained model and convolutional neural network

**DOI:** 10.1101/2024.01.28.577612

**Authors:** Yiqi Chen, Yang Qi, Yingfu Wu, Fuhao Zhang, Xingyu Liao, Xuequn Shang

**Affiliations:** School of Computer Science, Northwestern Polytechnical University, Xi’an, Shaanxi 710072, China; College of Information Engineering, Northwest A&F University, Yangling, Shaanxi 712100, China

**Keywords:** Transposable Elements, BERT, CNN, Hierarchical classification

## Abstract

Transposable Elements (TEs) are abundant repeat sequences found in living organisms. They play a pivotal role in biological evolution and gene regulation and are intimately linked to human diseases. Existing TE classification tools can classify classes, orders, and superfamilies concurrently, but they often struggle to effectively extract sequence features. This limitation frequently results in subpar classification results, especially in hierarchical classification. To tackle this problem, we introduced BERTE, a tool for TE hierarchical classification. BERTE encoded TE sequences into distinctive features that consisted of both attentional and cumulative *k-mer* frequency information. By leveraging the multi-head self-attention mechanism of the pre-trained BERT model, BERTE transformed sequences into attentional features. Additionally, we calculated multiple *k-mer* frequency vectors and concatenate them to form cumulative features. Following feature extraction, a parallel Convolutional Neural Network (CNN) model was employed as an efficient sequence classifier, capitalizing on its capability for high-dimensional feature transformation. We evaluated BERTE’s performance on filtered datasets collected from 12 eukaryotic databases. Experimental results demonstrated that BERTE could improve the F1-score at different levels by up to 21% compared to current state-of-the-art methods. Furthermore, the results indicated that not only could BERT better characterize TE sequences in feature extraction, but also that CNN was more efficient than other popular deep learning classifiers. In general, BERTE classifies TE sequences with greater precision. BERTE is available at https://github.com/yiqichen-2000/BERTE.

## 1. INTRODUCTION

A type of interspersed repeats called transposable elements (TEs) can undergo positional movement. They are found in almost all prokaryotes and eukaryotes, typically in large numbers. TEs not only constitute a large proportion of biological genomes, but also encode proteins as well as complex noncoding regulatory sequences, thus play an important role in gene regulation and genome evolution **(Wicker, *et al*., 2018)**. For the first time, Barbara McClintock discovered TEs in Zea mays **(McClintock, 1941)**. She found that some genes not only have the ability to relocate, but they can also be activated or deactivated in response to specific environmental conditions or during different stages of cellular development. In 1989, Finnegan proposed the first TE classification system, which included two categories **(Finnegan, 1989)**, RNA (class I or retrotransposons) or DNA (class II or DNA transposons), based on their transposition intermediate. Wicker *et al*. further standardized a hierarchical classification system for eukaryotic TEs **(Wicker, *et al*., 2007)**, including the levels of class, subclass, order, superfamily, family, and subfamily. For example, the class I in the highest level can be divided into several orders: *LTR, DIRS, PLE, LINE*, and *SINE*. The *LTR* can be further divided into five superfamilies: *Ty1/Copia, Ty3/Gypsy, Bel/Pao, Retroviruses, and Endogenous Retroviruses (ERV)*. Currently, the TE analysis of classification has become an extensively researched topic to provide deeper insight on transposable elements **(Goerner-Potvin and Bourque, 2018)**.

With the widespread popularity of NGS (next-generation sequencing) in decades **(Hu, *et al*., 2021)**, genomes are sequenced in large quantities. TE sequence repositories are well established in the context of the extensive development of TE annotation methods, such as Repbase **(Jurka, *et al*., 2005)**, Dfam **(Hubley, *et al*., 2016)**, PGSB Repeat Database **(Spannagl, *et al*., 2016).** Meanwhile, the evolution of Transposable Element (TE) classification tools is rapidly progressing, driven by the expansion of repositories and advancements in computing algorithms. Among traditional classification methods, RepeatModeler 1.0 uses RECON **(Bao and Eddy, 2002)**, RepeatScout **(Price, *et al*., 2005)**, TRF **(Benson, 1999)** and RMBlast to classify TEs based on similarity comparisons between consensus sequences and the ones from databases. REPCLASS **(Feschotte, *et al*., 2009)** utilizes three modules for classification: homology, structural features, and target site duplication. However, its classification capabilities are limited to the class and order levels, and it does not have the ability to classify at the superfamily level. LTR_retriever **(Ou and Jiang, 2018)** is a multithreading-empowered Perl program for *LTR* elements classification based on sequence structural identification. Specifically, these three methods use traditional classification strategies, which are time-consuming and have limited classification completeness for large datasets.

As artificial intelligence continues to develop in computer science, some popular machine learning and deep learning methods have found successful applications in the realm of bioinformatics **(Wu and Zhao, 2019)**. In terms of machine learning methods, TEclass **(Abrusan, *et al*., 2009)** uses LIBSVM **(Chang and Lin, 2011)** to classify repeats of different size categories. LTRdigest **(Steinbiss, *et al*., 2009)** uses local alignment-based and hidden Markov model-based algorithms for *LTR* element classification. PASTEC **(Hoede, *et al*., 2014)** utilizes HMM profiles to classify TE consensus sequences using TE structural feature and sequence similarity. TE-Learner **(Schietgat, *et al*., 2018)** is a framework that employs a random forest approach for the classification of *LTR* elements. On the other hand, TIR-learner **(Su, *et al*., 2019)** integrates a basic neural network with a homology-based method to classify *TIR* elements. TransposonUltimate **(Riehl, *et al*., 2022)** is a software specifically engineered for the classification, annotation, and detection of TEs. It selects relative *k-mer* frequencies and binary protein features as input data, which are subsequently processed by a random forest model for predictive analysis.

A recent machine learning method called Inpactor2 **(Orozco-Arias, *et al*., 2023)** classified *LTR* elements using Convolutional Neural Networks. Within these methods, only simple artificial intelligence algorithms were implemented, and they could not efficiently handle complex data from multiple databases. Meanwhile, most methods were only suitable for identifying certain TE elements, such as *TIR* or *LTR*. The rapid development of deep learning has brought numerous applications in various fields. Convolutional Neural Network (CNN) has been widely used in computer vision and natural language processing. This popularity stems from three fundamental advantages **(Gu, *et al*., 2018)**: equivalent representations, sparse interactions, and parameter sharing. These advantages not only streamline the training process by reducing the number of parameters but also bolster the model’s generalization performance. In the field of bioinformatics, the CNN model is widely applied in prediction, as demonstrated by **(Saman Booy, *et al*., 2022; Zeng, *et al*., 2016; Zhang, *et al*., 2022; Zhuang, *et al*., 2019)**. When it comes to TE classification, several previous methods been developed based on CNN and other deep learning methods. TERL **(da Cruz, *et al*., 2021)** applied a 2D CNN as a TE classifier after sequence pre-processing. The results showed that CNN has the potential to effectively predict TE sequences within the dataset compiled from seven databases. DeepTE **(Yan, *et al*., 2020)** combined CNN and HMMER3 **(Eddy, 2011)** to improve prediction performance, using *k-mer* frequency features of elements included in RepBase and PGSB repeat databases. Recently, TEclass2 **(Bickmann, *et al*., 2023)**, a deep learning method based on the Longformer **(Beltagy, *et al*., 2020)** architecture was proposed. It trained a single model to classify both order and superfamily categories meanwhile provided a simple web page for online inference with uploaded data. TEclass2 was trained on datasets from Repbase and Dfam, which filtered many existing TE categories after pre-processing. Despite utilizing a single model for TE classification, the evaluation metrics suggest that there is potential for further enhancement in its performance. The Longformer architecture inherently faces challenges in managing its attention mechanisms. These mechanisms may inadvertently focus on parts of the sequence that lack biological relevance, or they may display undesirable attention biases. In its default configuration, TEclass2 demands substantial graphical memory. The study demonstrates that TEclass2 operates on multiple GPUs, including an A100 with 80GB. This poses a challenge to replicate the process (training and inference) on regular servers. On the other hand, some research groups have utilized advanced natural language processing techniques to extract features from biological sequence data, such as a novel language representation model called BERT (bidirectional encoder representations from transformers) **(Devlin, *et al*., 2018)**. Current word embedding approachs used in NLP (natural language processing) can be applied to transform the contextual connections between amino acid or nucleotide sequences, which are treated as natural languages. Lee *et al*. proposed Bert-enhancer **(Le, *et al*., 2021)**, which converted enhancer sequences into a numerical matrix based on a pre-trained BERT model. After extracting features using BERT, Bert-enhancer feeds the features into a CNN model to predict enhancers. Charoenkwan *et al*. proposed BERT4Bitter **(Charoenkwan, *et al*., 2021)** to improve the prediction of bitter peptides. Followed by a BiLSTM classifier **(Siami-Namini, *et al*., 2019)**, the BERT model with 12 encoders directly transforms the original amino acid sequence without relying on any structural information.

From this idea, we propose a transfer learning-based classification method called BERTE. It uses a BERT pre-trained model and cumulative *k-mer* frequency vectors for feature extraction, and then uses a CNN classifier for TE hierarchical classification. The features and benefits of BERTE are reflected in the following aspects. First, BERTE uses a transfer learning method, employing a pre-trained BERT model for the purpose of feature extraction. Secondly, multiple *k-mer* frequencies, rather than a single one, are also selected as a type of input feature. Finally, both features are concatenated as the final input feature, which is then fed into a CNN model for classification. In comparison, the running time gap between all methods is not significant, but in *k-mer* countering, BERTE runs the fastest.

The implementation of Horner’s Rule **(Borodin, 1971)** achieves a notable speed increase through polynomial computation optimizations. It’s important to highlight that the dataset used in this study surpasses the size of those used in prior research, including 12 databases spanning various kingdoms. The experimental findings demonstrate that BERTE is capable of swiftly extracting sequence features and building a hierarchical model for effective TE classification. Overall, BERTE exhibits superior performance compared to existing methods for TE classification.

### Highlights

- **The first attention feature extraction module based on BERT transfer learning is proposed for high-precision TE hierarchical classification.** In this study, a BERT pre-trained model was used to extract attention features from TE sequences. Specifically, we took the DNA sequences in a certain length at both ends and fed them into the pre-trained model to obtain embedding features.
- **A *k-mer* frequency accumulation strategy is innovatively proposed to further enhance feature extraction.** As a second feature, we computed multiple *k-mer* frequencies of the TE sequence and concatenated these vectors as cumulative *k-mer* frequency features. Specifically, 4-mer, 5-mer and 6-mer frequencies were computed separately using Horner’s Rule.
- **BERTE adopts a top-down structure to achieve high-precision hierarchical classification of TEs.** For different TE categories (*i.e.*, class, order, and superfamily), we trained a separate model for each. In our study, there were 7 categories for TE, corresponding to 7 models. Unlike TEclass2, which is a large all-in-one classification method with a single model, BERTE achieved higher accuracy in most categories and required fewer computing resources.
- **The most comprehensive TE library to date has been built for training and testing of BERTE**. TE sequences used in this study were gathered from twelve public TE databases. Following this, we executed a series of pre-processing steps to prepare the final dataset for model training and testing. This dataset, comprising 932,944 sequences across 315 species, stands as the most extensive dataset for TE classification to date.
- **Compared to other methods, BERTE achieved best performance on imbalanced datasets.** In the currently accessible public datasets, it’s common to encounter imbalances within a single category. For instance, in our curated *LTR* training dataset, the sequence counts for *Bel-Pao* and *ERV* stood at 3,668 and 11,708 respectively, while those for *Copia* and *Gypsy* were significantly higher, with 150,651 and 210,498 sequences respectively. The F1-scores of BERTE on Bel-Pao and ERV are much higher than those of other methods.

## 2. MATERIALS AND METHODS

### 2.1 The overview of BERTE workflow

BERTE, as depicted in Figure 1, integrates two modules BERT and CNN, to facilitate a three-step hierarchical classification of TEs. Firstly, we obtained the pre-trained BERT model from Google **(Devlin, *et al*., 2018)** and truncated each sequence at both ends in a certain length leading to two sequence fragments. The two fragments were joint together. Secondly, we derived the 4-mer, 5-mer, and 6-mer sub-sequences from each joint fragment and inputted their respective frequencies into a pre-trained BERT model. We then extracted the output from the final encoder’s hidden layer and concatenated it with the corresponding vectors of the three sub-sequences (*i.e.*, the 4-mer, 5mer, and 6-mer frequency vectors of each joint fragment were concatenated). The final concatenated output was used as attentional features. Thirdly, we obtained the cumulative *k-mer* frequency vector for each full-length sequence. This vector is a concatenation of the frequency vectors for 4-mer, 5-mer, and 6-mer. We used this cumulative *k-mer* frequency as the second feature. Finally, both features were concatenated as the final sequence features. They were fed into the CNN classifier for training and prediction of different categories, enabling hierarchical TE classification.

**Figure 1.**
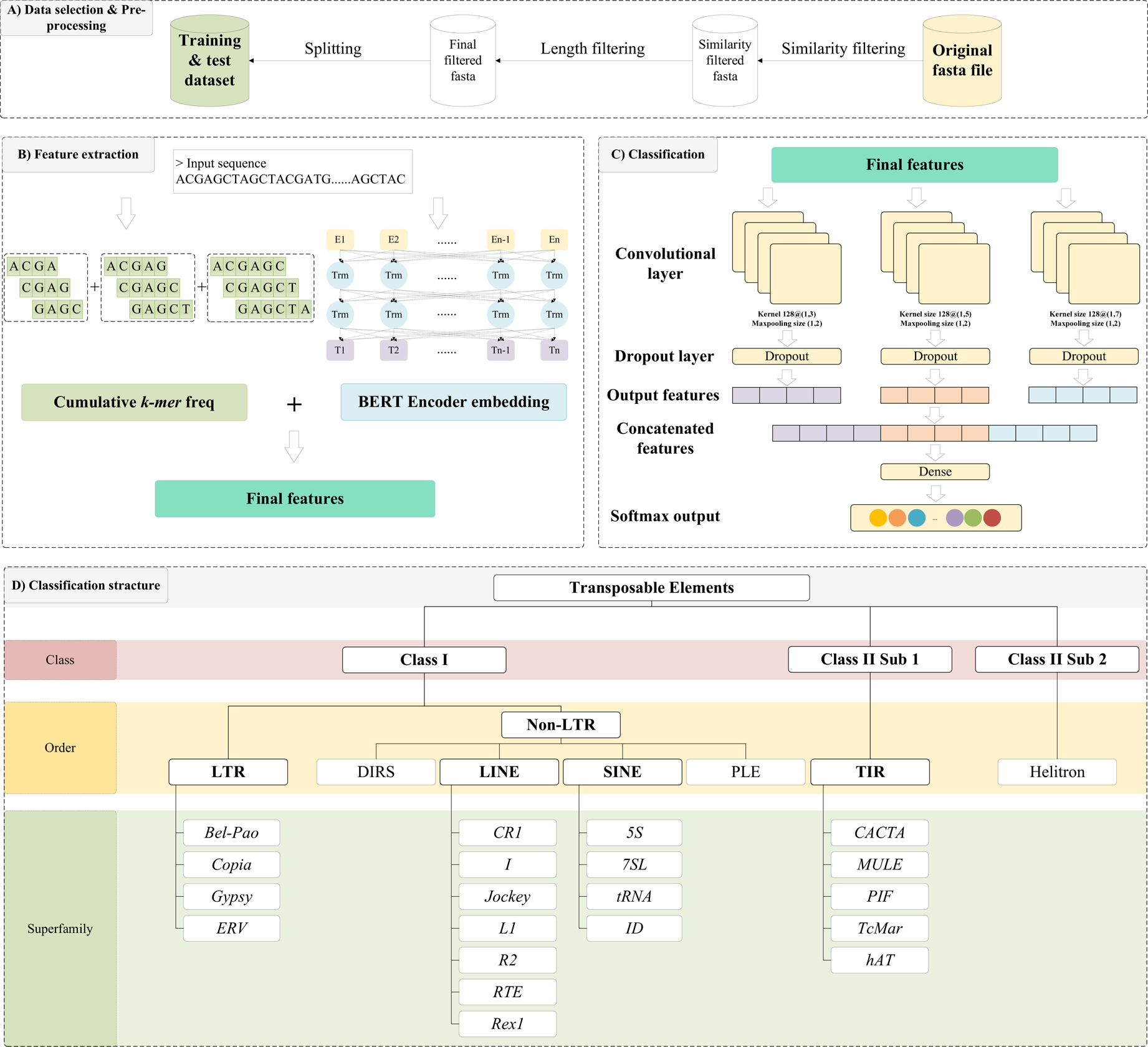
The overall workflow of the proposed method. A) Data selection and pre-processing. We collected sequences in fasta format from 12 databases and conducted filtration based on sequence similarity and length. Then, we split the data into training and testing datasets. B) Feature extraction. We transformed the dataset into two features. First, three types of *k-mer* frequencies were calculated separately and then concatenated as the cumulative *k-mer* frequency feature vector. Second, a pre-trained BERT model was used to extract attentional features. The two features were concatenated together as the final feature. C) The principle of Classification. We implemented a parallel CNN model to classify the input sequence features, using three convolutional layers with different kernel sizes. D) The hierarchical structure of TE classification. Unlike the popular hierarchical structure proposed in **(Wicker, *et al*., 2007)**, we adopted the remaining TE orders and superfamilies to build the classification structure after pre-processing.

#### 2.1.1 Feature extraction module based on BERT

BERT is a model that relies on transformer encoders (Vaswani, *et al*., 2017) and uses self-attention mechanisms, which are highly effective in NLP. The original BERT is divided into two steps: unsupervised pretraining on two tasks, masked-language modeling (MLM) and next sentence prediction (NSP), and then fine-tuning the pre-trained model for downstream tasks to get the final output of prediction probability (Devlin, *et al*., 2018).

In the BERT model, the input vector for each word consists of three embeddings: token embeddings, segment embeddings, and position embeddings. Token embedding is the representation of each word, which must be included in the vocabulary. Within this embedding, words need to be encoded by different segmentation methods. Segment embedding is used as a discriminator in the sentence pair. It distinguishes whether each word belongs to the first or second sentence. Position embedding aims to make BERT learn the sequential information of inputs. The position embedding encodes the relative and absolute positions of each word in the sentence, as well as the distance between different words in the sentence.

The BERT model architecture contains multiple identical transformer encoder blocks, which are connected back and forth to process the context using the attention mechanism and processes all the words simultaneously in the sentence. Each transformer encoder block contains two sub-layers: a multi-head attention layer and a feed-forward neural network (FFN) layer **(Bebis and Georgiopoulos, 1994)**, with residual connections and layer normalization between the sub-layers. Each multi-head attention layer contains multiple attention heads based on scaled dot product attention, allowing the attention mechanism to integrate various behaviors as knowledge using different subspace representations of queries, keys, and values. The formula is listed as follows:

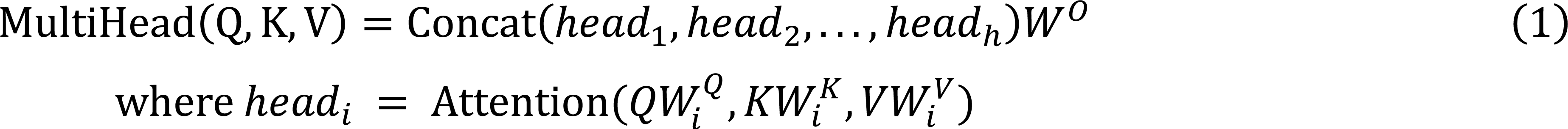

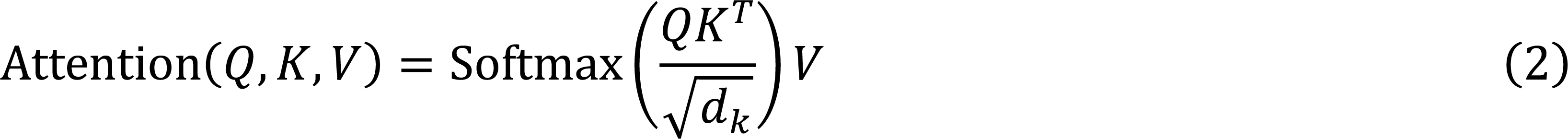

where 𝑄 (Query), 𝐾(Key) and 𝑉 (Value) are all derived from different linear transformations of input word embeddings to obtain information representations of different subspaces. 𝑑_𝑘_ is the dimension of 𝐾 and 𝑊^𝑄^_*i*_, 𝑊^𝐾^_*i*_, 𝑖 𝑖 𝑊^𝑉^_*i*_ and 𝑊^𝑂^_*i*_ are weight matrices. The FFN layer consists of two linear transformations with rectified linear unit 𝑖 𝑖 (ReLU) activation function in the middle. The formula is listed as follows:

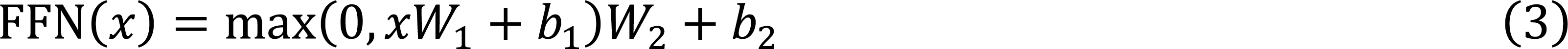

In this study, the sequences were fed into a pre-trained BERT model for inference. Firstly, due to the varying sequence length in the dataset, the fixed-length fragments at both ends were truncated, corresponding to the cumulative *k-mer* sub-sequences. In this step, fragments of 1,000, 1,250, and 1,500 base pairs (bp) were truncated from the left and right ends, respectively. These truncated fragments were then concatenated, resulting in total lengths of 2,000, 2,500, and 3,000 bp. After that, the concatenated fragments from both ends are converted into token sentences according to the vocabulary list. In this study, we took 500 tokens to represent each sequence, along with three special tokens, resulting in a total of 503 tokens in total after segmentation (*i.e.*, sequence tokens + a classifier token [CLS] and two separator [SEP] tokens). The selected BERT model contains 4 encoders, and the [CLS] token in the last hidden layer state of the fourth encoder is extracted for each *k-mer* sub-sequence. Then, we concatenated them as the final BERT feature vectors.

Since only the [CLS] token embeddings were extracted, the loss function in this module is focused on the sentence-level task in pretraining. The formula is listed as follows:

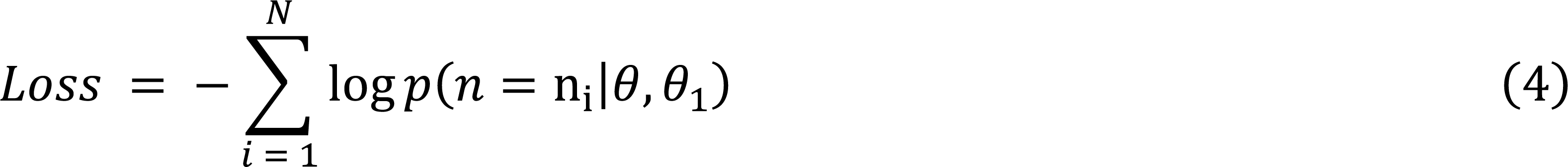

where 𝑛 represents the sentence pair and 𝑛_𝑖_ represents whether the sentence pair is consecutive or not. 𝑁 represents the number of sentence pair. Notably, 𝜃 represents the parameters in BERT Encoder, and 𝜃_1_ represents the sentence-level prediction task parameters in BERT Encoder.

#### 2.1.2 Feature enhancement module based on cumulative *k-mer* frequency

*k-mer* refers to a consecutive sequence of 𝑘 nucleotides. For example, a 2-mer can be “AT”, “CG” and so on. Given a fix 𝑘, there are 𝑙 − 𝑘 + 𝑠 *k-mer*s generated from a sequence of length 𝑙, where 𝑠 is the sliding window size. The *k-mer* frequency refers to the number of occurrences of each distinct *k-mer* in a given sequence.

In this study, a polynomial computation method called Horner’s Rule (Borodin, 1971) is used for *k-mer* frequency counting, which improves the computational speed. Each nucleotide in the *k-mer* subsequence (A, C, G, and T) is encoded as a number (0, 1, 2, and 3). The digital sequence is considered a quadratic number in base-4 and then converted to a decimal number in base-10, which indicates the index in the array. Horner’s rule is used to efficiently convert numbers from quaternion to decimal representation. For example, it converts a 5-mer ACCTG to 01132 (in base-4) and then to 94 (in base-10), mapping it to the 94^th^ cell in the array. Note that the length of the array storing all *k-mer*s is 4^k^. Horner’s Rule is calculated as follows:

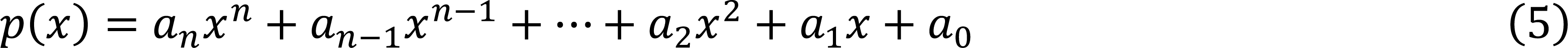

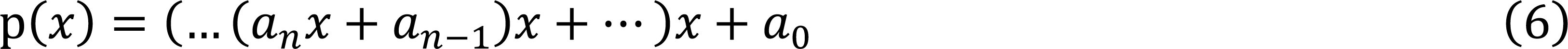

where 𝑎^0^, 𝑎^1^, …, 𝑎^𝑛^, and 𝑥 are known real numbers. Horner’s Rule converts (5) into the (6). The computational complexity of finding the value of each term in (5) and then summing is 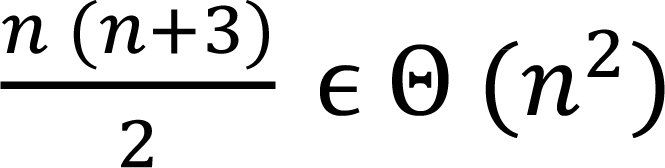. After applying Horner’s Rule, it only needs to perform 𝑛 multiplications to obtain the value of a polynomial of order 𝑛. Therefore, the complexity is Θ(𝑛) in (6), which brings a significant improvement compared to solving the first polynomial directly in (5).

#### 2.1.3 CNN classification module

After feature extraction, the features are fed into the CNN model for classification. CNN is a powerful neural network designed for processing image data **(Gu, *et al*., 2018)**. In recent bioinformatics research, CNN has been also used to study the information of topology structure **(Sun, *et al*., 2020)**, dinucleotide one-hot encoder **(Lv, *et al*., 2021)**, PSSM profiles **(Le, *et al*., 2020)** (image-like data with fewer pixels), or even word embedding **(Le, *et al*., 2019)**.

A CNN is made up of these main types of layers: input layer, convolutional layer, activation layer, pooling layer, and fully connected layer. By stacking these layers on top of each other in a specific order, a complete convolutional neural network can be constructed. The convolutional layer, which is the cornerstone of a CNN, utilizes a convolutional kernel to perform feature transformations. This process is responsible for most computations within the network. The pooling layer performs downsampling and ensures translation invariance. The fully connected layer serves to map the learned distributed feature representation to the sample label space, acting as a ’classifier’ in the entire CNN. In our CNN model, we adopted a parallel strategy for CNN architecture construction. At the same time, the input passed through three Conv2D layers activated by ReLU functions and three max pooling layers first. These three Conv2D layers had different kernel sizes, which were (1,3), (1,5), and (1,7). Following each max pooling layer, a dropout layer was applied. The outputs from these dropout layers were then concatenated into a single vector, which was subsequently fed into a fully connected layer. Finally, we used Softmax function **(Heaton, 2017)** for classification. It maps the outputs of the neural network to a range between 0 and 1, transforming them into probabilities corresponding to each class label, thereby facilitating classification. The convolution operation is formulated as follows:

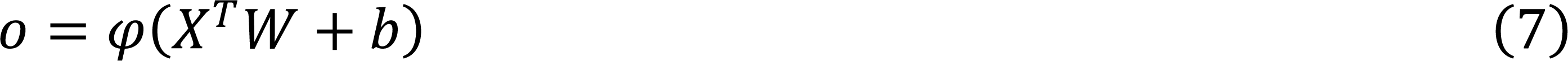

where 𝑜 is the output of the convolution, 𝑋 denotes the region of the convolution kernel, 𝑊 and 𝑏 correspond to its weight and bias, and 𝜑 is the activation function. In this study, we chose ReLU =max(𝑥, 0) as the activation function for the neural network. The Softmax function, used for multiclass classification, converts the outputs into probability values ranging between 0 and 1, indicating the likelihood of each possible class. The Softmax formula is listed as follows:

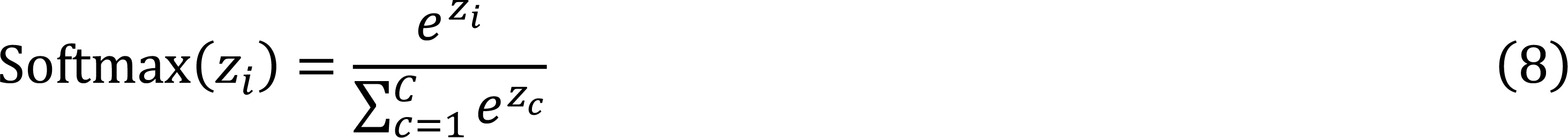

where 𝑧_𝑖_ is the output value of the 𝑖^th^ node and 𝐶 is the number of output nodes, *i.e.*, the number of categories in classification.

Cross-entropy (CE) (de Boer, *et al*., 2005) was chosen as the loss function for the model, meanwhile the Adam algorithm (Kingma and Ba, 2014) was chosen as the optimizer. The formula is listed as follows:

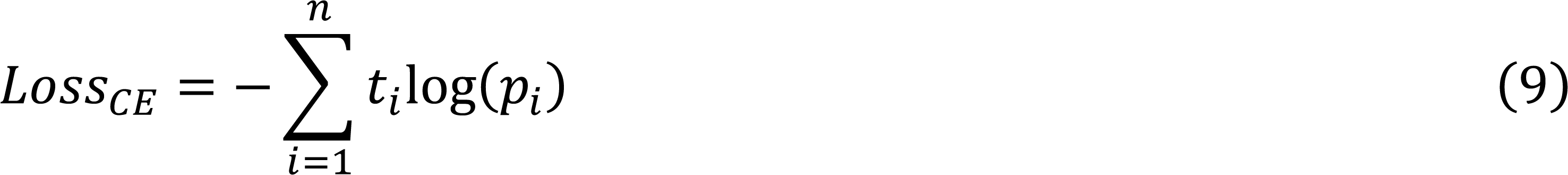

where 𝑡_𝑖_ is the truth label and 𝑝_𝑖_ is the Softmax probability for the 𝑖^th^ class. We used Python 3.8 and Tensorflow **(Abadi, *et al*., 2016)** to build the model.

### 2.2 Datasets pre-processing and evaluation metrics

#### 2.2.1 Datasets

Datasets from twelve TE repositories were selected in this study, covering plants, animals, and fungi (Table 1). In detail, we collected a total of 932,944 TE sequences from various databases, including APTEdb **(Pedro, *et al*., 2021)**, CicerSpTEdb **(Mokhtar, *et al*., 2023)**, ConTEdb **(Yi, *et al*., 2018)**, DPTEdb **(Li, *et al*., 2016)**, Dfam **(Hubley, *et al*., 2016)**, FishTEDB **(Shao, *et al*., 2018)**, MnTEdb **(Ma, *et al*., 2015)**, PGSB **(Spannagl, *et al*., 2016)**, Repbase **(Jurka, *et al*., 2005)**, RepetDB **(Amselem, *et al*., 2019),** RiTEdb **(Copetti, *et al*., 2015)**, SoyTEdb **(Du, *et al*., 2010)** and TREP **(Wicker, *et al*., 2002)**. After that, we filtered sequences during pre-processing, resulting in the retention of representative superfamilies only, as shown in Table 2. In the model training process, the dataset was split into training, validation, and testing sets in an 8:1:1 ratio.

**Table 1.**
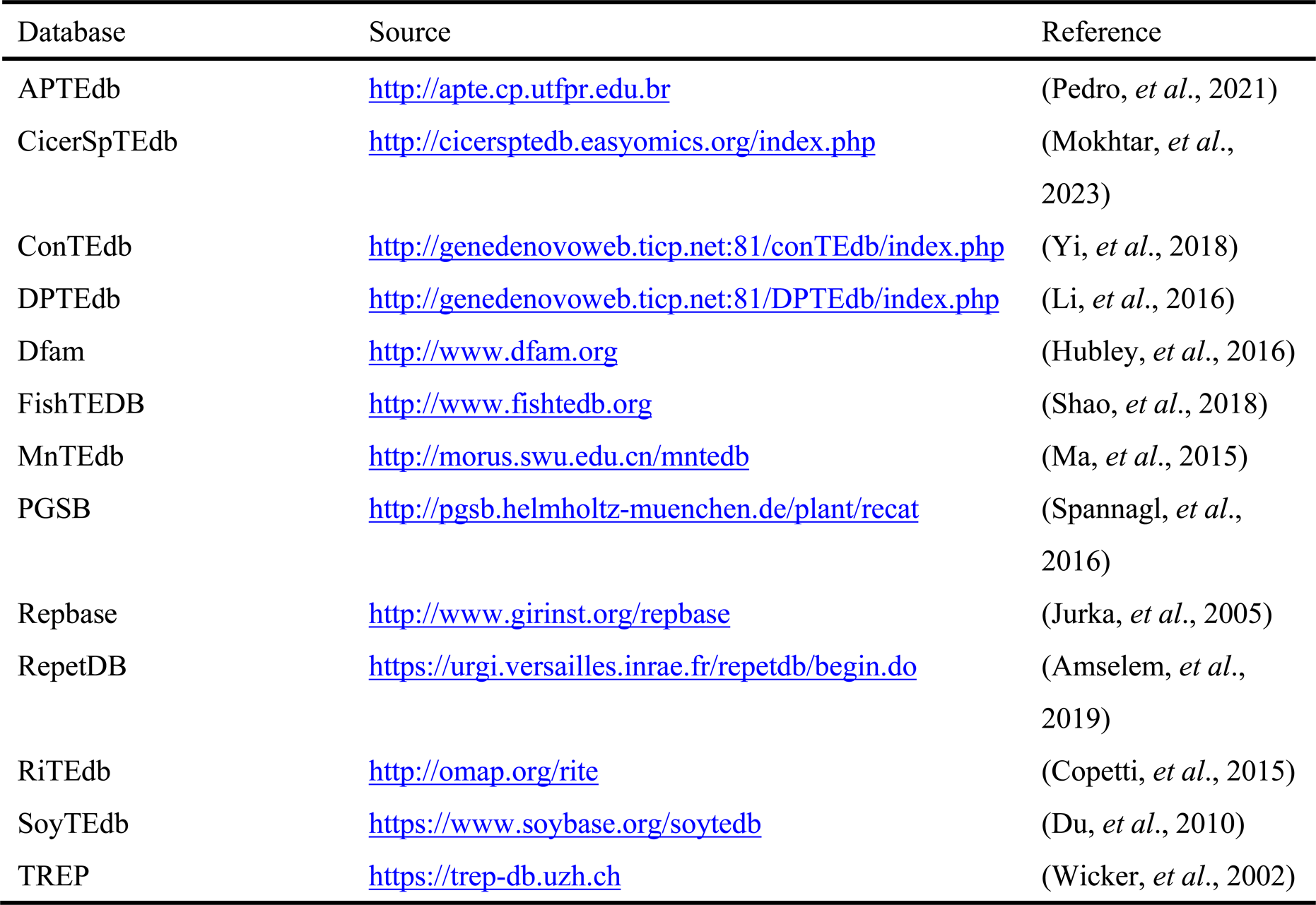
The source of databases used in this study.

**Table 2.**
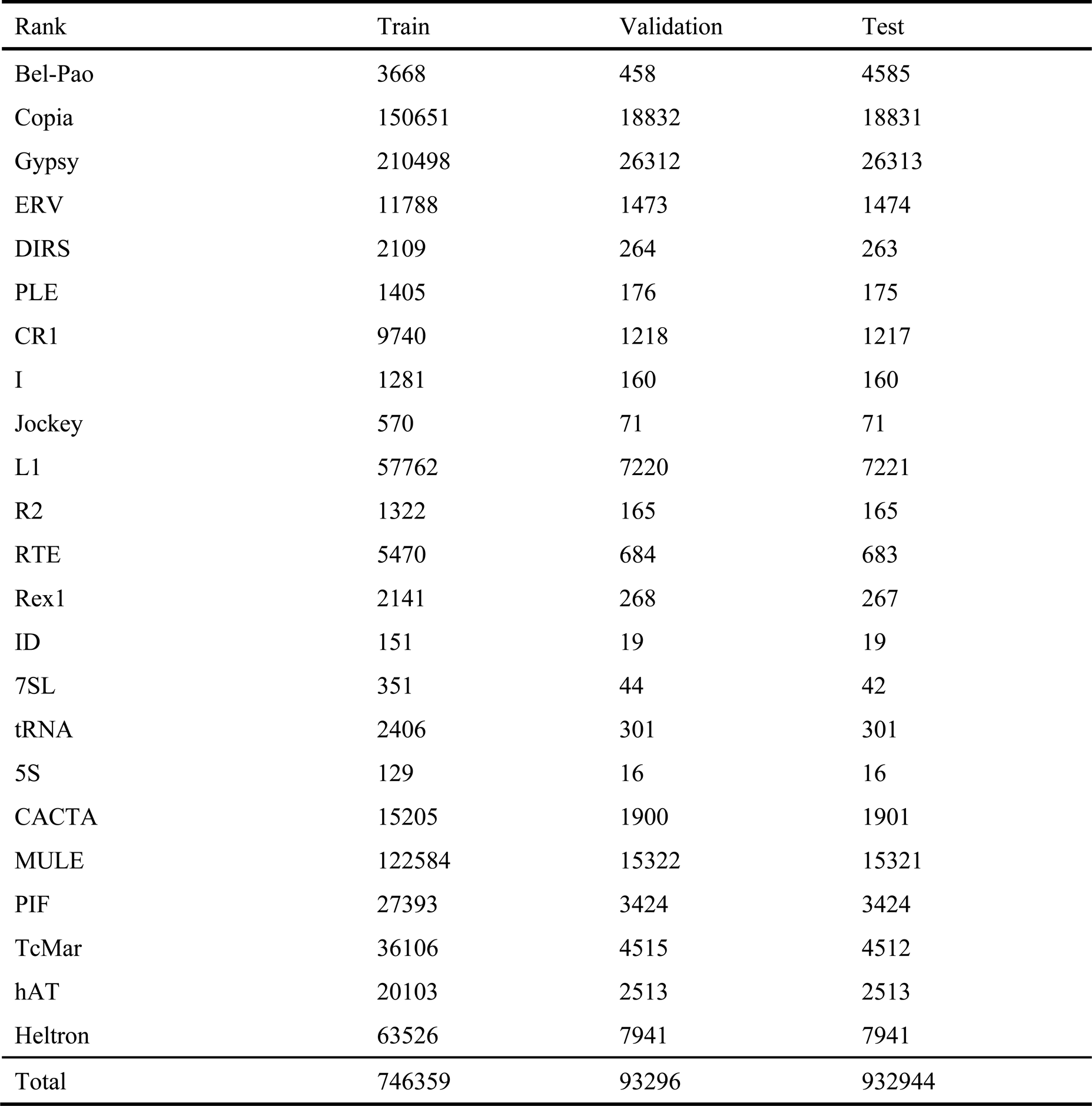
The sequences numbers in different orders and superfamilies. The dataset had been divided into training, validation and testing ones.

| Rank | Train | Validation | Test |
| --- | --- | --- | --- |
| Bel-Pao | 3668 | 458 | 4585 |
| Copia | 150651 | 18832 | 18831 |
| Gypsy | 210498 | 26312 | 26313 |
| ERV | 11788 | 1473 | 1474 |
| DIRS | 2109 | 264 | 263 |
| PLE | 1405 | 176 | 175 |
| CR1 | 9740 | 1218 | 1217 |
| I | 1281 | 160 | 160 |
| Jockey | 570 | 71 | 71 |
| L1 | 57762 | 7220 | 7221 |
| R2 | 1322 | 165 | 165 |
| RTE | 5470 | 684 | 683 |
| Rex1 | 2141 | 268 | 267 |
| ID | 151 | 19 | 19 |
| 7SL | 351 | 44 | 42 |
| tRNA | 2406 | 301 | 301 |
| 5S | 129 | 16 | 16 |
| CACTA | 15205 | 1900 | 1901 |
| MULE | 122584 | 15322 | 15321 |
| PIF | 27393 | 3424 | 3424 |
| TcMar | 36106 | 4515 | 4512 |
| hAT | 20103 | 2513 | 2513 |
| Heltron | 63526 | 7941 | 7941 |
| Total | 746359 | 93296 | 932944 |

#### 2.2.2 Pre-processing

In order to minimize data imbalance and redundancy, several pre-processing steps for the data are conducted as follows: (i) removing duplicate sequences based on 100% similarity, which preliminarily reduces redundancy, and (ii) sequences which are less than 100 bp in length were removed. Besides, only sequences that include “A”, “C”, “G”, and “T” at least once each are retained. (iii) Among the remaining sequences, those with unknown labels and without labels are filtered out to ensure that the final dataset contains specific label information. After the pre-processing steps mentioned above, the filtered dataset was divided into training sets, validation, and test sets according to the training strategy. Thus, the final dataset contains 932,944 TE sequences, with 746,359, 93,296, and 93,289 sequences corresponding to the training, validation, and test sets, respectively.

#### 2.2.3 Evaluation metrics

To evaluate the performance of classification, we calculated four different measurement metrics: recall, precision, accuracy, and F1-score. These four parameters are defined as follows:

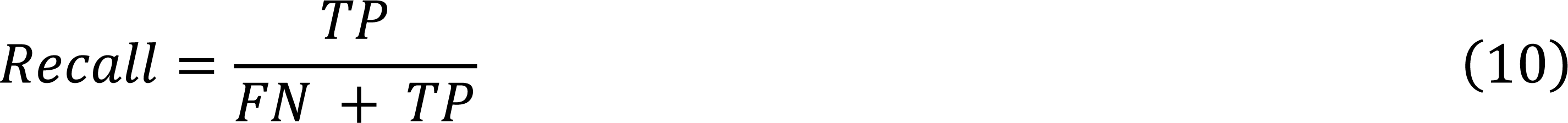

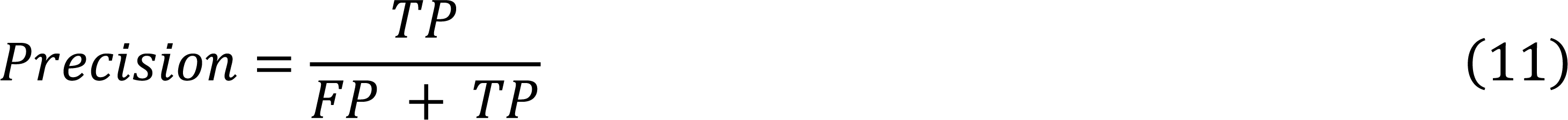

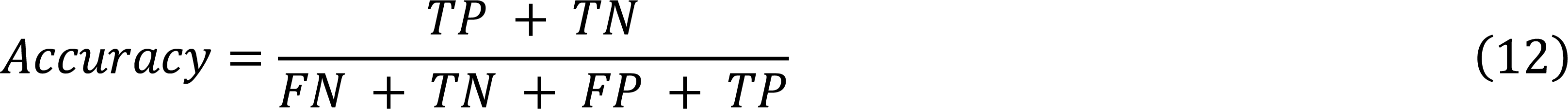

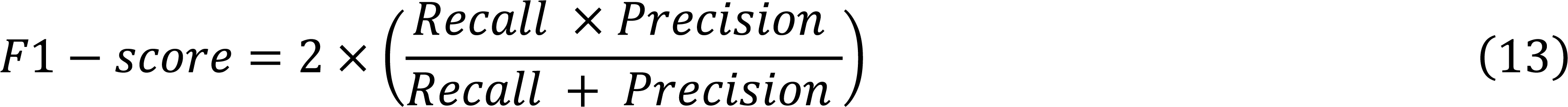

where TP, TN, FP and FN denote true positives, true negatives, false positives and false negative, respectively.

## 3. RESULTS

### 3.1 BERTE improves the performance of TE classification

To evaluate the effectiveness of BERTE in TE classification, we used the split testing dataset for analysis. Sequence features were extracted by a pre-trained BERT model and cumulative *k-mer* frequency, and subsequent CNN model accepts these features for training and prediction to classify TE sequences. In 10-fold cross-validation, BERTE trained different models for each category within a tiered system, classifying sequences from top to bottom. As for the other two methods, we chose their default parameters for training and testing.

Within method comparison, the evaluation metrics include accuracy, precision, recall, and F1-score. By analyzing the evaluation metrics across various methods, we can ascertain the performance of each classification technique. As shown in Tables 3-4 and Figure 2, the evaluation results showed that BERTE outperforms both DeepTE and TERL, both DeepTE and TERL are in default settings. The accuracies of these three methods are all above 60%, with BERTE and DeepTE achieving 90% in most models. In terms of the rest metrics, TERL behaved the worst, showing a large gap compared to the other two methods. It is probably because the pre-processing strategy of TERL introduces many noise features, which uses one-hot encoding and pads feature vectors to ensure equal length. Compared to DeepTE, BERTE performed better overall, and it has obtained a large lead in *LTR* and *LINE*. In the *LTR* category, BERTE outperformed DeepTE with an F1-score of 79% compared to DeepTE’s 71%. Similarly, in the Non-LTR category, BERTE again led with an F1-score of 83%, while DeepTE trailed at 75%. In conclusion, BERTE demonstrates the best results across most categories, indicating outstanding classification performance. As mentioned above, BERTE obtained higher F1-scores in *LTR* and *LINE* categories, which indicates its ability to cope with imbalanced data. In the *LTR* category, the F1-scores of *Bel-Pao* and *ERV* are 55% and 81%, respectively, significantly surpassing DeepTE’s scores of 34% and 77%. As shown in Tables 3-4, the numbers of *Bel-Pao* and *ERV* are much less than *Copia* and *Gypsy*, leading to a serious imbalance issue. However, there is room for improvement in BERTE’s handling of data imbalances. The evaluation metrics indicate a significant drop-off for minority class data compared to the majority class, suggesting the need for further optimization.

**Table 3.** The BERTE’s F1-score performance compared with state-of-the-art methods in different models on class categories. For each model, we present both the overall macro-average F1-score and the F1-scores for each individual rank.

| Model | Category | F1-score |  |  |
| --- | --- | --- | --- | --- |
|  |  | TERL | DeepTE | BERTE |
| ClassI |  | 0.62 | 0.90 | 0.93 |
|  | LTR | 0.90 | 0.97 | 0.98 |
|  | Non-LTR | 0.12 | 0.83 | 0.88 |
| ClassII |  | 0.81 | 0.88 | 0.90 |
|  | Sub1 | 0.89 | 0.95 | 0.96 |
|  | Sub2 | 0.33 | 0.81 | 0.84 |

**Table 4.** The BERTE’s F1-score performance compared with state-of-the-art methods in different models on order categories. For each model, we present both the overall macro-average F1-score and the F1-scores for each individual rank.

| Model | Category | F1-score |  |  |
| --- | --- | --- | --- | --- |
|  |  | TERL | DeepTE | BERTE |
| LTR |  | 0.24 | 0.71 | 0.79 |
|  | Bel-Pao | 0.00 | 0.34 | 0.55 |
|  | Copia | 0.58 | 0.83 | 0.88 |
|  | Gypsy | 0.67 | 0.89 | 0.91 |
|  | ERV | 0.41 | 0.77 | 0.81 |
| Non-LTR |  | 0.34 | 0.76 | 0.80 |
|  | DIRS | 0.00 | 0.71 | 0.75 |
|  | PLE | 0.00 | 0.56 | 0.66 |
|  | LINE | 0.96 | 0.98 | 0.98 |
|  | SINE | 0.14 | 0.80 | 0.81 |
| LINE |  | 0.22 | 0.75 | 0.83 |
|  | CR1 | 0.12 | 0.82 | 0.87 |
|  | I | 0.00 | 0.52 | 0.69 |
|  | Jockey | 0.00 | 0.50 | 0.73 |
|  | L1 | 0.86 | 0.96 | 0.97 |
|  | R2 | 0.10 | 0.77 | 0.84 |
|  | RTE | 0.00 | 0.80 | 0.84 |
|  | Rex1 | 0.01 | 0.84 | 0.87 |
| SINE |  | 0.44 | 0.82 | 0.81 |
|  | 7SL | 0.44 | 0.91 | 0.93 |
|  | tRNA | 0.91 | 0.96 | 0.97 |
|  | 5S | 0.00 | 0.69 | 0.64 |
|  | ID | 0.10 | 0.71 | 0.69 |
| TIR |  | 0.48 | 0.78 | 0.82 |
|  | CACTA | 0.21 | 0.55 | 0.67 |
|  | MULE | 0.77 | 0.93 | 0.93 |
|  | PIF | 0.44 | 0.80 | 0.82 |
|  | TcMar | 0.50 | 0.87 | 0.89 |
|  | hAT | 0.41 | 0.73 | 0.78 |

**Figure 2.**
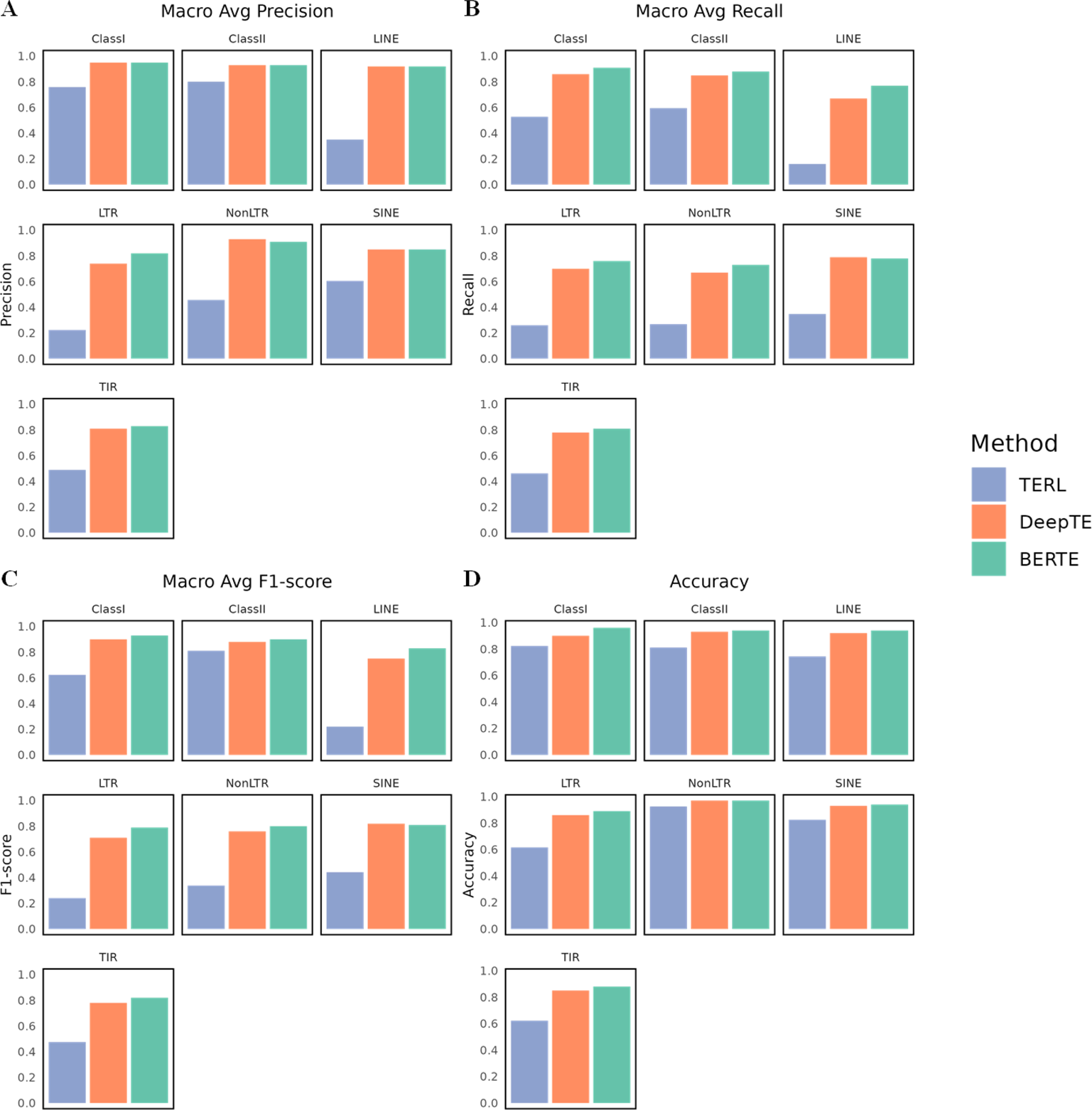
Comparison of the proposed method’s performance with state-of-the-art methods in different evaluation metrics. A, B, C, D corresponds to macro average precision performance, macro average recall performance, macro average F1-score performance and accuracy performance separately.

### 3.2 Hyperparameter optimization

A crucial aspect of training deep learning models is the optimization hyperparameters, which assists in achieving the optimal performance **(Feurer and Hutter, 2019)**. In this study, we used a combination of BERT and CNN for TE classification, so the optimal hyperparameters for each model need to be determined individually.

First, we conducted hyperparameter optimization for the BERT model. Google has published pre-trained BERT models (Devlin, *et al*., 2018) at multiple scales whose feature representation performance may vary. In this study, we fed the training set to the pre-trained models for fine-tuning, and 10-fold cross-validation was used to determine the optimal hyperparameters. As shown in Table 5, several typical BERT models were selected for fine-tuning: namely BERT-Tiny (with 4.4 million parameters, 2 transformer layers, and 128 hidden embedding sizes), BERT-Mini (with 11.3 million parameters, 4 transformer layers, and 256 hidden embedding sizes), BERT-Small (with 29.5 million parameters, 4 transformer layers, and 256 hidden embedding sizes), BERT-Medium (with 41.7 million parameters, 8 transformer layers, and 512 hidden embedding sizes) and BERT-Base (with 101.1 million parameters, 12 transformer layers, and 768 hidden embedding sizes).

**Table 5.**
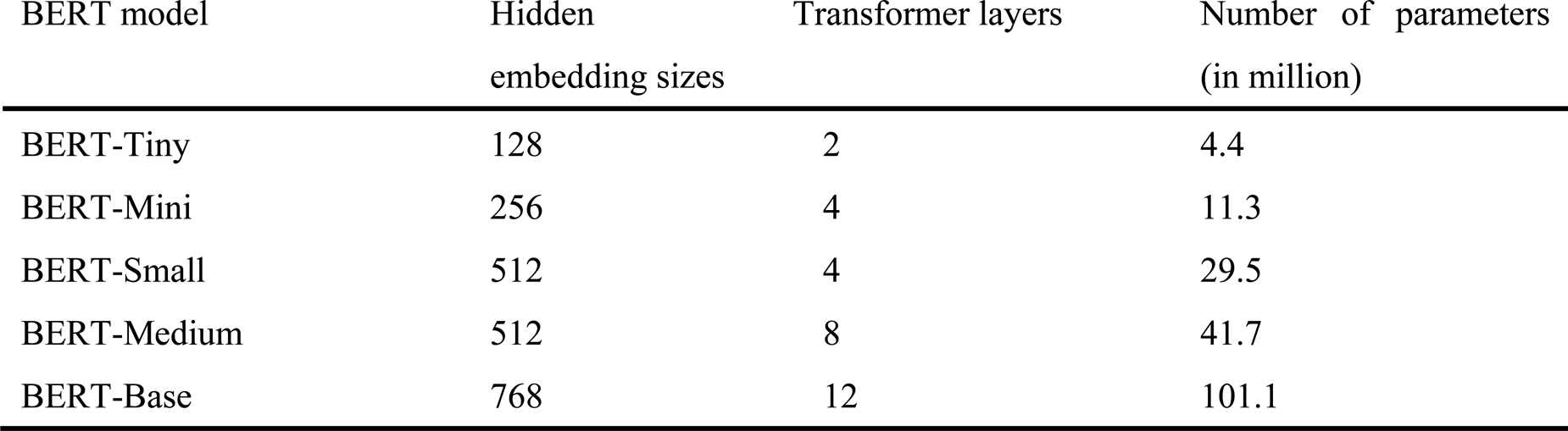
Different parameters for each pre-trained BERT model. Ultimately, BERT-Mini was selected as our tool of choice for extracting attentional features.

The hyperparameters of the BERT model consists of batch size, maximum input sequence length, learning rate, and training steps. We tried several combinations of these hyperparameters to obtain the optimal parameters. The results showed that the best evaluation metrics was obtained from the BERT-Mini model. Since BERT-Mini has a relatively small model size and number of parameters, and also runs the second fastest, we chose it as the optimal feature representation for these pre-trained BERT models.

Secondly, we optimized the hyperparameters for the CNN model. The hyperparameters of our CNN model include the number of convolutional kernels, batch size, and epochs. We conducted 10-fold cross-validation and tried several combinations to determine the optimal parameters. It is found that the optimal results are obtained when the number of convolutional kernels is set to (1,3), (1,5), (1,7) for each convolutional layer, the number of batch sizes is 64, and the number of epochs is 50. The final CNN hyperparameters are shown in Table 6.

**Table 6.** The hyperparameters in our parallel CNN module.

| CNN Hyperparameters | Values |
| --- | --- |
| Epoch | Default 50, with EarlyStopping |
| Batch size | 64 |
| Learning rate | 1e-7 in minimum, warming up to 1e-4 |
| Kernel size | (1,3), (1,5), (1,7) separately for each Conv layer |

Note: We have set the default epoch to 50 and enabled EarlyStopping in TensorFlow to mitigate overfitting. Subsequently, we implemented the warmup strategy (Gotmare, *et al.*, 2018) during model training. The Learning rate is initially increased linearly from a minimum of 1e-7 to 1e-4, after which it is gradually decreased to 1e-7. Regarding the Kernel size, we implemented three distinct sizes for the three convolutional layers: (1,3), (1,5), (1,7) respectively.

### 3.3 Comparison to traditional biological features and NLP features

In order to validate the effectiveness of BERT-encoded features in classification, we compared BERT with other common traditional bioinformatics and NLP features. In this study, we regenerated these features based on our TIR dataset, and compared them with our proposed features using a CNN classifier. Common bioinformatics features such as *k-mer* frequency, pseudo-dinucleotide composition (PseDNC) **(Chen, *et al*., 2013)**, and pseudo k-tuple nucleotide composition (PseKNC) **(Chen, *et al*., 2015)** were used in many studies. Additionally, three common NLP models used in bioinformatics analysis, fastText **(Bojanowski, *et al*., 2017)**, Word2Vec **(Mikolov, *et al*., 2013)** and ELMo **(Sarzynska-Wawer, *et al*., 2021)**, can also be employed to encode features. The fastText software was used directly to train on our dataset. We used Word2Vec from gensim Python package **(Rehurek and Sojka, 2011)** to train on our dataset. ELMo has pre-trained models, so we re-trained the pre-trained ELMo model, extracted features, and passed them into the subsequent classifier.

Based on 10-fold cross-validation, we compared the feature extraction results of BERT with the other six methods mentioned above, along with the final hybrid features used in our study. The hybrid features outperformed in all metrics, as delineated in **Supplementary Table S1**.

### 3.4 Models based on different classifiers

Based on the mixture of pre-trained BERT-Mini and cumulative *k-mer* features, we fed them into different classifiers including, CNN, GRU (Chung, *et al*., 2014) and BiLSTM to evaluate the performance of different classifiers. We optimized the hyperparameters for each classifier by experimenting with various combinations of hyperparameters. Table 7 lists the TIR sequences prediction performance of different classifiers based on the optimal hyperparameter combinations, in 10-fold cross-validation. The result shows that the CNN-based classification is the most effective. The CNN classifier obtained in all evaluation metrics, and notably run the fastest among all methods. It can be seen that the CNN model was able to classify the TIR sequence most effectively both in performance and time consumption. Therefore, we adopt the CNN model as the final classifier in BERTE.

**Table 7.**
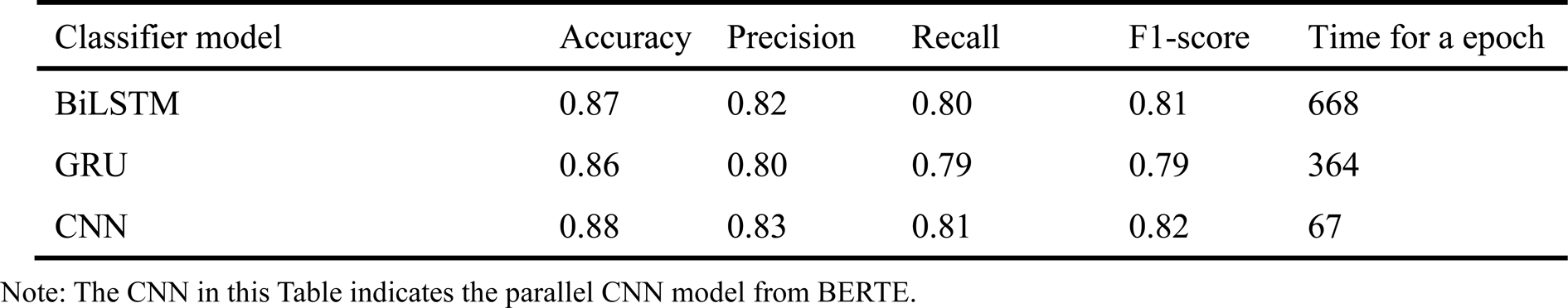
Performance evaluation on the representative TIR sequences using different deep learning classifiers, based on 10-fold validation.

### 3.5 BERTE uses attention to better classify TE sequence

To analyze the learning effectiveness of BERT, we extracted the features of each encoder layer using the pre-trained BERT-Mini model on the TIR dataset. The features extracted by the last BERT layer improve the ability to distinguish different TEs. This shows that BERT can learn informative feature representations from input sequences in higher level, as can be seen in **Supplementary Figure S1**.

To further analyze how the BERT model learns sequence information, we extracted and analyzed the attentional weights from this model. For each sequence, the attentional weights from different heads in different layers were analyzed. Since the [CLS] token embeddings in the last layer are further used for classification, we computed the [CLS] attentional weights from the other tokens. Specifically, the attentional weights from the other tokens for [CLS] in different heads were combined in each layer. As a result, we obtained a vector of values at each layer that corresponds to the overall [CLS] attention weights from the other tokens. The attentional weights are shown in **Supplementary Figure S2**.

## 4. DISCUSSION

This study demonstrates that the classification accuracy of TE sequences can be significantly improved by leveraging the multi-head self-attention mechanism of BERT and the high-dimensional feature transformation capability of CNN. Through the overall experiments, BERTE proved to be an effective tool for TE sequence analysis, excelling in feature extraction. In the method comparison, the results indicate that BERTE is the most useful and outperforms other methods across most metrics. It has been observed that the combination of traditional *k-mer* frequency features and BERT encoding features yields better classification results than using BERT encoding features alone.

The outstanding performance of BERTE can be attributed to its effective feature extraction capability as well as its powerful classifier. DNA sequences can be treated like natural language. By employing the self-attention mechanism, the model can process contextual information and parallelize all words in the input sentence. This capability enhances its efficiency in handling TE sequences of diverse lengths. Recent deep learning-based methods use traditional bioinformatic features such as *k-mer*, PseDNC, PseKNC, *etc*. However, these methods exhibit limitations when it comes to portraying the relationships between sequences that occur before and after. BERTE employs a more advanced self-attention mechanism to comprehend the contextual information of the sequence, while also ensuring its ability to handle variable-length sequences, making it highly suitable for the analysis of TE sequences. When comparing BERTE with other methods, it can be found that BERTE performs better across most TE categories. This can be attributed to the hybrid features of the self-attention module and cumulative *k-mer* frequency, as well as the excellent classification ability of the CNN. In terms of feature extraction, the hybrid features from BERT and *k-mer* provide more effective sequence information. Secondly, in terms of classifiers, CNN achieved the highest scores across all evaluation metrics, demonstrating excellent prediction ability. It should be noted that all existing methods for TE classification cannot handle imbalanced data effectively. Imbalanced data is a common occurrence in the currently available TE databases, and the classification of TEs remains a challenge under situations with large numerical gaps. From the experimental results, it is evident that BERTE has the most comprehensive classification performance, likely due to its use of more practical sequence representations.

## 5. CONCLUSION

The method BERTE introduced in this study facilitates the high-precision hierarchical classification of TE sequences. BERTE utilized the concatenated TE features consisted of multi-head self-attention information from BERT and cumulative *k-mer* frequency information. By using pre-trained BERT model, semantic and syntactic information extracted from natural language can be transferred to analyze nucleic acid sequence data, as BERT is still a state-of-the-art method used in various NLP tasks. After feature extraction, BERTE adopted a parallel CNN as a classifier, leveraging its capability for high-dimensional feature transformation. Consequently, this study suggests that employing BERT-based embeddings with CNN provides a more accurate approach to transfer biological language. Notably, it is an effective attempt to implement attention mechanism in TE analysis, which can be visualized through computing attention weights. From the experimental results, the BERT-based features can represent TE sequences more efficiently, providing an application to treat DNA sequences as natural languages. The model proposed in this study could also be implemented to analyze other biological sequences after making certain adjustments.

### Data availability

The source code of BERTE is available at https://github.com/yiqichen-2000/BERTE.

## Author contributions

X.Q.S. supervised the project. Y.Q.C., X.Q.S., and X.Y.L. conceived and designed this work. Y.Q.C., X.Y.L., and Y.Q. conceived, designed, and implemented the BERTE pipeline. Y.Q.C., and Y.Q. preprocessed the data and coordinated data release. Y.Q.C., X.Y.L., and F.H.Z. analyzed the performance of algorithms developed in this study. Y.Q.C., and X.Y.L. wrote the paper. All authors have read and approved the final version of this paper.

